# Magnesium hydroxide nanoneedles derived from *Anthocleista schweinfurthii* Gilg (Loganiaceae) support mesenchymal stromal cell proliferation and wound healing

**DOI:** 10.1101/2024.04.29.591621

**Authors:** Francois Eya’ane Meva, Rita Pereira, Sandrine Elodie Ngnihamye, Tchangou Njiemou Armel Florian, Agnes Antoinette Ntoumba, jean Batiste Hzounda Fokou, Thi Hai Yen Beglau, Marcus N. A. Fetzer, Marilyn Kaul, Bianca Schlierf, Ulrich Armel Mintang Fongang, Phillipe Belle Ebanda Kedi, Simone Veronique Fannang, Marietta Herrmann, Christoph Janiak

**Author notes:** Corresponding author: Francois Eya’ane Meva.

## Abstract

Multiple metallic nanoparticles are able to promote cellular and tissue health, but these nanoparticles can be difficult to synthesize and can also cause unintended side-effects. Here, we study the effects on wounds healing and bone reparation of Mg(OH)_2_ from *Anthocleista schweinfurthii* Gilg (Loganiaceae) leaves (AS), which are local to the Africa region and have been used in traditional medicine to treat injuries. Mg(OH)_2_ nanoneedles were synthesized from aqueous extracts of *Anthocleista schweinfurthii* Gilg (Loganiaceae) leaves (AS) and magnesium nitrate. The quick polydispersing and crystallized Mg(OH)_2_-metal interface was found to be covered in plant secondary metabolites. We call this compound Mg(OH)_2_-AS. Using an acute dermal toxicity experiment on animal model, we determined that Mg(OH)_2_-AS is safe for topical application. *In vitro* experiments suggest that Mg(OH)_2_-AS has anti-inflammatory potential, and *in vivo* wound healing assays in Wistar rats indicate that Mg(OH)_2_-AS can enhance wound healing. To investigate Mg(OH)_2_-AS effects on the cellular level, we used bone marrow mesenchymal stromal cells (BM-MSCs). In contrast to pure Mg(OH)_2_ or AS, cell viability and proliferation were not impaired by Mg(OH)_2_-AS. Cell morphology remained unchanged upon media supplementation with Mg(OH)_2_-AS. Preliminary results further indicate enhanced osteogenic differentiation of BM-MSCs in media supplied with ascorbic acid, β-glycerophosphate and dexamethasone and addition of Mg(OH)_2_-AS. These findings motivate further research towards the inclusion of the material in implants for bone fracture healing.

## Introduction

Musculoskeletal disorders, which include more than 150 different conditions, affect around 1.71 billion people and are the leading cause of disability. Musculoskeletal disorders reduce mobility and dexterity, negatively impacting society [1]. Fractures, caused by traumatic injury, are a major factor in disability, resulting in lost work and income and an increased societal economic burden. Living tissue responds to a traumatic event with mechanical failure, followed by an acute inflammatory reaction. The local inflammatory response involves an immediate and prolonged release of inflammatory mediators, that may be released in granules (histamine) or synthesized de novo (prostaglandins, cytokines), and plasma-derived, to initiate the healing response [2]. Likewise, protein denaturation is associated with inflammation and leads to various inflammatory diseases including arthritis [3]. Simple fractures can be treated simply with mechanical stabilization, whereas large segmental defects require more complex interventions like bone grafting [4]. Bone grafting has several drawbacks, including donor side morbidity and pain, motivating a search for alternative strategies to promote healing.

Bone marrow contains mesenchymal stromal cells (BM-MSCs) that can both self-renew and differentiate into skeletal tissues, including bone, cartilage and adipose tissue [5]. Moreover, their immunosuppressive ability establishes MSCs as a candidate cell type in tissue engineering, regenerative medicine and cell therapy for immune disorders [6]. Previous studies have shown that ceramic and metallic nanoparticles (hydroxyapatite, silica, silver, and calcium carbonate) can affect differentiation of MSCs into osteoblasts and adipocytes [7]. For example, Poly(D,L-lactic-co-glycolic acid; PLGA)/ Magnesium hydroxide (Mg(OH)_2_) nanoparticle scaffold in PLGA support chondrogenic healing of rat osteochondral defect sites *in vivo* [8]. This effect of Mg(OH)_2_ on MSC function and bone healing may be related to the fact that magnesium is known to seed calcium phosphate and subsequent growth of the bone tissue mineral component hydroxyapatite [9], determine intracellular protein and DNA synthesis, and induce osteogenic and osteoblastic differentiation [10].

Magnesium-based biomaterials (MBs) have been used as orthopedic implants for more than 100 years due to their desirable mechanical and osteopromotive properties [11]. However, current Mg-based materials have also negative aspects due to corrosion [12]. Metal-based nanoparticles are widely used in biomedicine due to their easy handling and functionalization, as well as their biocompatibility and improved physicochemical properties. In *in vivo* wound healing assay on adult mice, Mg(OH)_2_ biocomposites resulted in 1 cm^2^ burned wounds healing faster compared to control [13]. Bioceramics and Mg-based bioglasses that have been extensively tested as degrading bone replacement materials are stepwise substituted by newly formed bone tissue [10]. Moreover, magnesium hydroxide-incorporated PLGA composite scaffolds have been suggested as efficient delivery system for bone morphogenic protein (BMP)2, with osteoinductive and anti-inflammatory effects *in vitro* and *in vivo* [14].

Nanoparticles such as Mg(OH)_2_ can be synthesized using biogenic methods including the use of plants and microorganisms. During the biological synthesis of nanoparticles, the enzymes, active metabolites, or phytochemicals of the biotic source work as reducing and capping agents, thus allowing the generation of the particles [15]. By controlling the physico-chemical properties of the nanoparticles, the capping molecules can produce a synergy of action, with the latter, who can transport them into the target cells as a nanocarrier, thereby enhancing activity [16]. For example, metallic nanoparticles of gold modified with 11-mercaptoundecanoic acid were effective in enhancing adipogenic differentiation and weakening osteogenic differentiation in bone marrow mesenchymal stromal cells (BM-MSCs) from Sprague-Dawley rats due to generating higher levels of reactive oxygens species (ROS) [17]. Similarly, metallic silver nanoparticles and ions have been shown to stimulate adipogenic and osteogenic differentiation of human MSCs in a concentration dependent manner, while no effect was seen on chondrogenic differentiation after 21 d of incubation [18]. It is possible that Mg nanoparticle function can be similarly enhanced by synthesizing those nanoparticles from biogenic sources.

Leaf decoctions of *Anthocleista schweinfurthii* Gilg (Loganiaceae) have been traditionally used for wound healing as in the treatment of fever. *In vitro* assays have confirmed their antibacterial, and 2,2-diphenyl-1-picrylhydrazyl (DPPH) radical scavenging potential [19]. Two xanthones derivatives: 1,8-dihydroxy-2,6-dimethoxyxanthone and 1-hydroxy-3,7,8-trimethoxyxanthone known, respectively, as swertiaperenin and decussatin have been extracted from the leaves [20]. Decoction of bark and leaves of *A. schweinfurthii* have been used for the treatment of pain, injury and inflammatory diseases [21–23]. Here, we describe the effect of characterized *Anthocleista schweinfurthii* (AS) mediated magnesium hydroxide nanoneedles on wound healing *in vivo* and on bone marrow mesenchymal stromal cells *in vitro*. We characterize the material and consider its potential for wound healing and regenerative therapies, e.g. to support bone formation.

### Experimental

#### Collection, authentication, and preparation of extract

*Anthocleista schweinfurthii* Gilg (Loganiaceae) leaves (Figure 1) were harvested in the Faculty of Medicine and Pharmaceutical Sciences Botanical Garden (N04°11.585; E009° 39.585’), Littoral region, Cameroon, and authenticated at the Cameroon National Herbarium compared to a voucher specimen previously deposited number 32389/SRFCam. The freshly harvested leaves were washed with tap water and then with distilled water to remove all contaminants. They were then finely cut into 2 mm wide pieces to increase the contact surface with the solvent. 20 g of leaves were added to a beaker containing 100 mL of preheated distilled water (80 °C), and the mixture was kept under constant stirring for 5 minutes using a thermostatically-controlled hot plate equipped with a magnetic stirrer. After cooling to room temperature, the solution was filtered through Whatman n°1 paper and concentrated by removing residual solvent in an oven at 40 °C. Finally, the dry extract obtained was weighed to assess the extractable content determined by the following formula 1.

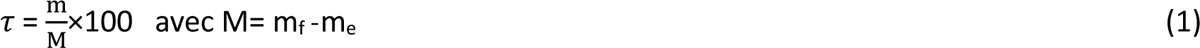

**Figure 1.**
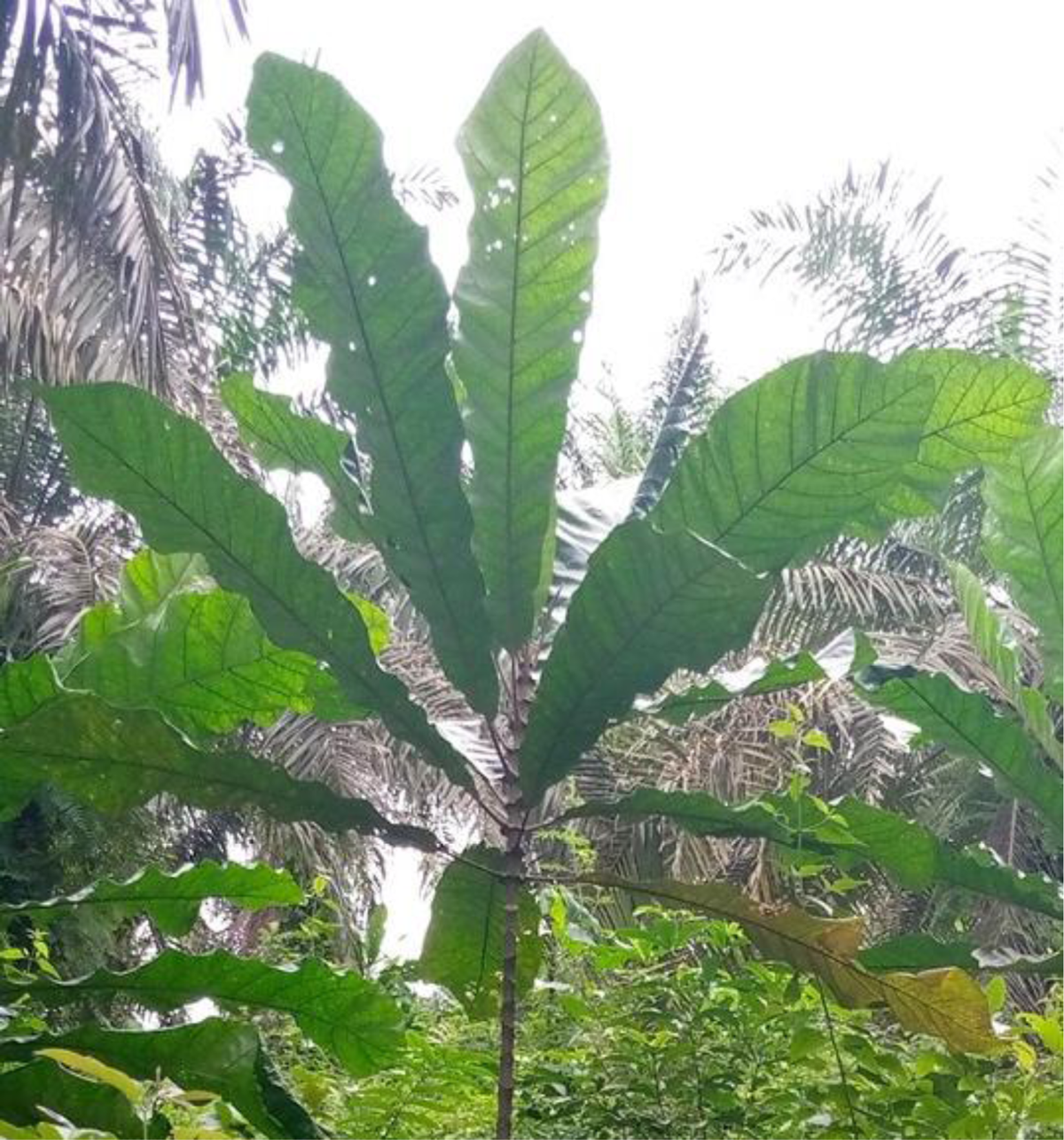
*Anthocleista schweinfurthii* (AS) in its natural habitat

τ: extractable content in percent (%); m: mass of dry extract (g);

M: mass of dry leaves (g); m_f_: mass of fresh leaves (g);

m_e_: mass of water contained in fresh leaves (g)

#### Phytochemical screening

The phytochemical screening to detect alkaloids, flavonoids, terpenoids, coumarins, saponins, anthocyanins, and anthaquinones was performed using le Thi et al established protocol [24].

#### Biological synthesis of Mg(OH)_2_ nanoparticles

For synthesis, 50 mL of 0.1 M magnesium nitrate hexahydrate salt Mg(NO_3_)_2_·6H_2_O (Roth) were preheated to 30, 50 and 70 °C. Then 10 mL of aqueous AS extract was added dropwise. The reaction medium was made basic by adding 1 mL of 5M KOH (VWR) and the mixture was kept under constant stirring for 2.5 h. After cooling the preparation to room temperature, 1 mL was taken to confirm nanoparticle formation by UV spectrophotometric scanning between 200 and 800 nm and by dynamic light scattering. The rest of the preparation was centrifuged at 6500 rpm (Hettich D-7200 Tuttlingen, Germany) for 30 minutes. The supernatant was replaced once with distilled water, and twice with 70% ethanol. The residue was then oven-dried at 60 °C under vacuum for 24 h and used for characterization studies. Control Mg(OH)_2_ nanoparticles were obtained using a modified protocol of Fellner et al, 2001 [26]. The precipitation of Mg(OH)_2_ was obtained by treating Mg(NO_3_)_2_ 6H_2_O by KOH in hot distilled water (80 °C). Mg(OH)_2_ solution was centrifuged and the precipitate was washed with distilled water and dried at 100 °C to obtain a white powder.

#### Mg(OH)_2_-AS solution preparation for wound healing assay

Centrifuged Mg(OH)_2_-AS nanoparticles were suspended in water and diluted to the desired concentration then applied to the wounds.

#### Mg(OH)_2_-AS solution preparation for MTT assay

Solutions were prepared following Escheverry-Rendon and co-workers method with modifications [27]. Concentration of Mg(OH)_2_-AS 1000 μg/mL and controls were prepared in 99.9% ethanol. The nanoparticle solutions were ultrasonicated before use. The ethanol solution was added to obtain the desired concentration of 150 and 300 μg/mL and evaporated under UV light for 30 minutes. The nanoparticles were suspended for 3-(4,5-Dimethyl-2-thiazolyl)-2,5-diphenyl-2H-tetrazolium bromide (MTT) tests and differentiation studies.

#### Ultraviolet-visible characterization (UV-Vis)

Ultraviolet-Visible spectroscopy was monitored using an aliquot of 2 mL of suspension on a P9 double beam spectrophotometer from VWR. Measurements were made between 200 and 800 nm.

#### Dynamic Light Scattering (DLS)

DLS was performed on a Malvern Nano S Zetasizer HeNe laser wavelength at 633 nm, from nanoparticles in water. Three runs were made per measurement.

#### Fourier-transformed infrared spectroscopy (FTIR)

Infrared spectroscopy was conducted using a Bruker Tensor 37 with attenuated total reflection, ATR unit by scanning between 600-4000 cm^-1^.

#### Powder X-ray diffraction (PXRD)

The PXRD diffraction of the nanoparticles was performed using a Bruker D2 Phaser powder diffractometer (Cu K-Alpha1 [Å] 1.54060, K-Alpha2 [Å] 1.54443, K-Beta [Å] 1.39225) by preparing a thin film on a low-background silicon sample holder.

#### Scanning electron microscopy (SEM) and energy-dispersive X-ray spectroscopy (EDX) investigations

Scanning electron microscopy (SEM) was carried out with a Jeol JSM−6510LV QSEM Advanced electron microscope with a LaB_6_ cathode at 20 kV equipped with a Bruker Xflash 410 (Bruker AXS, Karlsruhe, Germany) silicon drift detector for energy-dispersive X-ray spectrometric (EDX) analysis. The nanoparticle samples were sputtered with gold by using a JEOL JFC-1200 Fine Coater.

#### Transmission electron microscopy (TEM)

Electron micrographs of the nanoparticles were recorded using a JEOL JEM-2100 Transmission Electron Microscope at 200 kV with a TVIPS F416 camera system. The images were analyzed using Fiji software for particle size measurements.

#### Animal and ethical considerations

Healthy adult female Wistar albino rats weighing between 150 - 200 g were obtained from the Laboratory Animal Centre of the Department of Pharmaceutical Science, Faculty of Medicine and Pharmaceutical Science, University of Douala, Cameroon. The animals were sorted randomly in standard polypropylene cages in groups of three and maintained under standard conditions of temperature (24 ± 2 °C) and light (approximately 12 h/12 h light/dark cycle) with free access to standard laboratory diet and tap water *ad libitum*.

#### Acute dermal toxicity study

An acute dermal toxicity study in Wistar albino rats was performed in accordance with the OECD Guideline number 402 [28]. Groups were distributed according to the protocol in Banerjee et al, 2013 [29]. A total of 6 females Wistar albino rats were divided into two groups; one group was treated with 2000 mg/kg of Mg(OH)_2_-AS (dispersed in water and applied topically), and the other group were water was applied was untreated (the control group). One day prior to treatment, each animal was weighed, anaesthetized (intraperitoneal injection of both ketamine chlorhydrate (50mg/ml) with acepromazine (10 mg/kg)) prior to its back hair being clipped with an electric hair clipper. Each rat was then caged individually and left undisturbed for 24 hours. On the treatment day, the nanoparticles suspension was applied evenly to the exposed skin. The animals were checked twice daily for 14 days for signs of irritation, changes in behaviour, and possible mortality. Body weight measurement was taken daily for 14 days.

#### Wounds excision

Wound induction and monitoring was performed according to Nko’o et al 2020 [30]. Animals were anesthetized with intraperitoneal injection of ketamine (50 mg/kg) and acepromazine (10 mg/kg) 1 minutes prior to wounding to be done in 30 minutes. Prior to wounding, the area was prepared with 70% alcohol, the dorsal fur was clipped using an electric clipper, and the anticipated wound region was marked on the dorsal thoracic region (1 cm away from vertebral column and 5 cm away from the ear) of the back using a stainless steel stencil. Toothed forceps, a scaple, and pointed scissors were used to create full thickness excision wounds (1 cm^2^ area and 2 mm depth). The wound was blotted with cotton soaked in normal saline, then left uncovered and exposed. Animals were housed in separate cages, with the bedding changed daily. All treatments were given 24 h after wound creation as described below. Wound area, wound contraction, and epithelialization period were evaluated during the entire healing process.

##### Animal treatment

The treatment was performed following Nko’o et al 2020 with modifications [30]. Before the experiments, rats were given one week to acclimatize to laboratory conditions. The experimental rats were randomly divided into six groups of five rats per group. Rats in group I received no treatment (negative control). Rats in group II were treated with trolamine (0.67 %) ointment (positive control). Rats in groups III-V were treated with Mg(OH)_2_-AS suspension 25, 50, or 100 mg/kg, respectively, and rats group VI were treated with the plant extract AS. Treatment was gently applied topically: wounds were blotted with cotton swabs soaked in each suspension, then the wounded area was covered until reaching complete healing once daily. These treatments were performed during the light phase of the nyctemeral cycle and animals were used only once.

Determination of wound healing rate: The wound closure rate was determined as described by Shetty *et al.,* 2012 with modifications [31]. Briefly, the wound area of each rat was measured on days 1, 3, 5, 7, 9, and 11 after creation of the wound using semi-transparent tracing paper and a permanent marker. The tracing paper was positioned on a 1 mm^2^ graph sheet and traced out. The percent wound closure was calculated based on the measured wound areas using equation (2) below. The epithelialization time was determined as the number of days required after wounding for the scab to fall off, leaving no raw wounds behind.

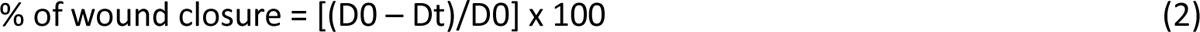

Where D0 is the wound size on day 0, and Dt is the wound size on day t.

##### Protein denaturation assay

As a hallmark of inflammation, inhibition of heat-induced denaturation of bovine serum albumin (BSA) was tested following Nikolaidis and Moschakis, 2017 with slight modifications [32]. The reaction medium consisted of 0.200 µL BSA (SIGMA Aldrich), 0.4%, w/v in PBS buffer, 2800 µL phosphate-buffered saline (pH 7.4) and 200 µL of nanoparticle solutions at various concentrations (200, 100, 50, 25 µg/mL). Distilled water served as a negative control and diclofenac was used as positive control, both tested under the same conditions as the nanocomposites. The mixture was first incubated at 37°C for 15 min and followed by incubation at 70 °C for 5 min [33]. After homogenization, the absorbance was measured at 660 nm. The percentage inhibition is calculated by eq. (3):

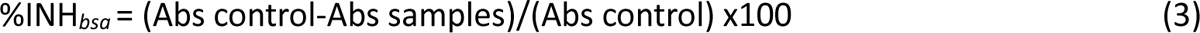

Abs: absorbance; % INH*_bsa_*: percentage inhibition.

### BM-MSC isolation and expansion

Mesenchymal stromal cells (MSCs) were isolated from bone marrow (BM) of two female donors of 82 and 83 years of age. BM was obtained as surgical waste material from acetabular reaming of patients receiving total hip replacement, with informed consent from the patient (ethical approval (187/18), University Hospital Würzburg, Germany). MSCs were isolated and expanded according to Pereira et al. [34]. Briefly, mononuclear cells were enriched from BM by Ficoll (Histopaque®-1077, Sigma-Aldrich, Germany) density gradient centrifugation (150 RCF for 5 min, Heraeus Multifuge X1R Centrifuge, Thermo Fisher Scientific, Germany). The cells were seeded at a density of 60000 cells/cm^2^ and expanded in culture medium (DMEM/F-12 GlutaMAX, Gibco, Germany) with the addition of 10% fetal calf serum (FCS, Bio&Sell, Germany), 1% Pen/Strep (Gibco), 1% HEPES (Sigma-Aldrich), and 5 ng/mL fibroblast growth factor (FGF, 100-18C, PeproTech, Germany) in a humidified atmosphere at 37 °C and 5% CO_2_. Culture medium was replaced three times a week. For passaging, cells were detached from culture vessels by trypsinization (T4174, Sigma-Aldrich), centrifuged, and washed, and the number of viable cells was determined with the trypan-blue dye (93595, Sigma-Aldrich) exclusion test. Cells were reseeded at a density of 20000 cells/cm^2^ or subjected for MTT tests or differentiation assays at passages 2-4.

### 3-(4,5-dimethylthiazol-2-yl)-2,5-diphenyl tetrazolium bromide (MTT) assay

The protocol described by Pereira and co-workers was used with few modifications [34]. For the MTT assay evaluation of metabolic activity of cells, MSCs of each donor were seeded in 96-well plates at a density of 10 × 10^3^ cells/well and treated the following day with nanoparticles at 150 and 300 μg/mL dosages in Dulbecco’s Modified Eagle Medium DMEM/F-12 (1:1) 1X, 1% GlutaMAX, 10% Foetal Calf Serum (FCS), 1% Penicillin/Streptomycin (P/S), 1% (4-(2-hydroxyethyl) piperazine-1-ethanesulfonic acid) (HEPES) media solutions. Basal cell culture media, Mg(OH)_2_, and dry AS ethanol extract were used as controls. After 24, 72 and 120 hours, the medium was removed and 10 μL of MTT solution (5 mg/mL) was added to the media in each well and incubated at 37 °C for 3 h. Formazan crystals resulting from cleavage of MTT were dissolved in 100 μL DMSO for 5 min with shaking and incubated 10 minutes. Each plate was read immediately in a microplate reader (Thermo Scientific, Waltham, MA) at a wavelength of 540 nm. Experiments were performed in duplicate. Cell metabolic activity is expressed in Figure 2 as percentage of the metabolic activity of untreated cells incubated in bsal media at each respective day, presented as the mean ± standard error of the mean, and compared using two-way ANOVA.

**Figure 2.**
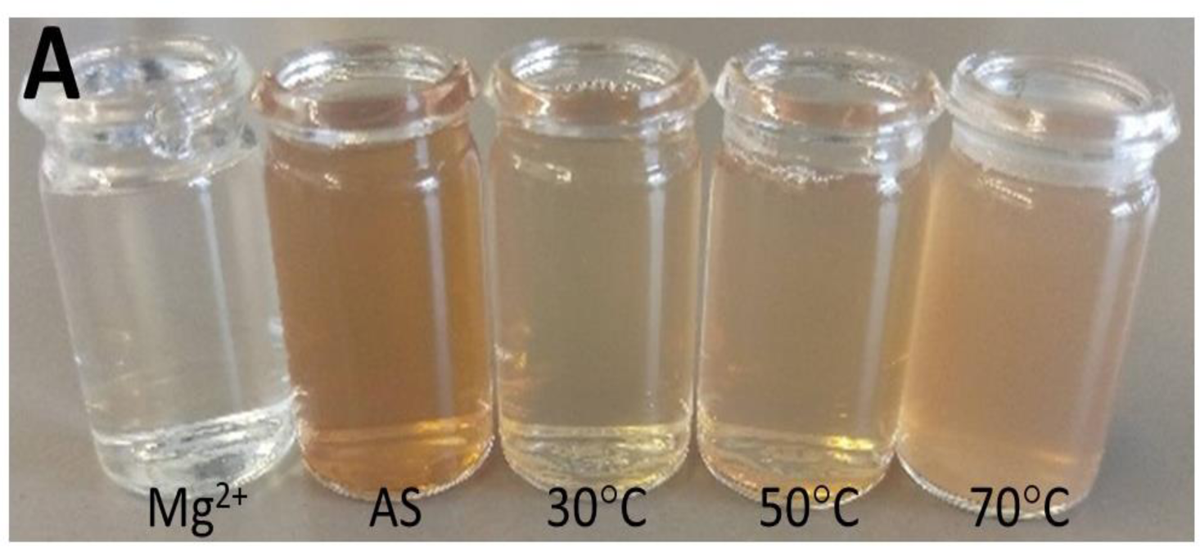
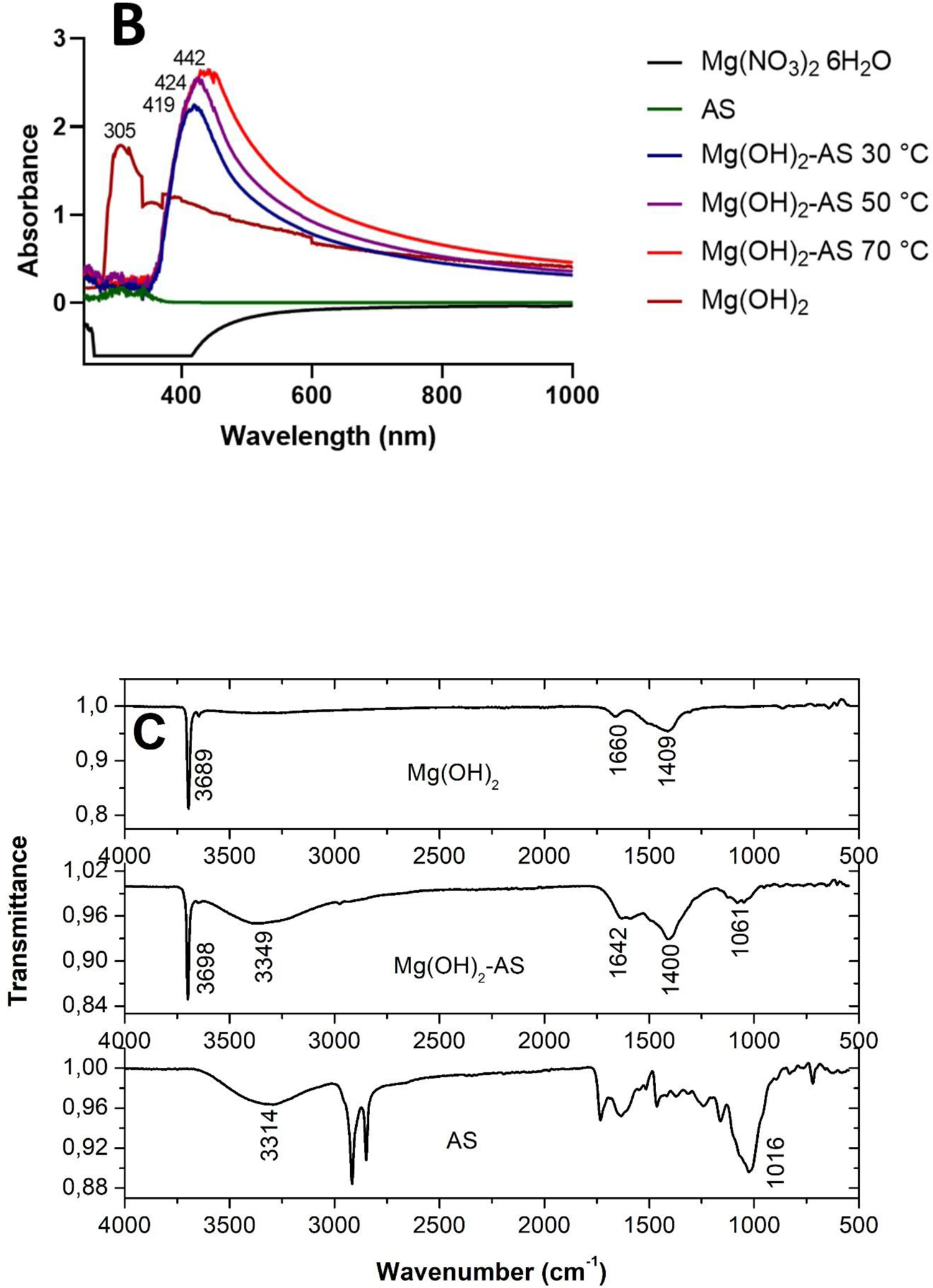
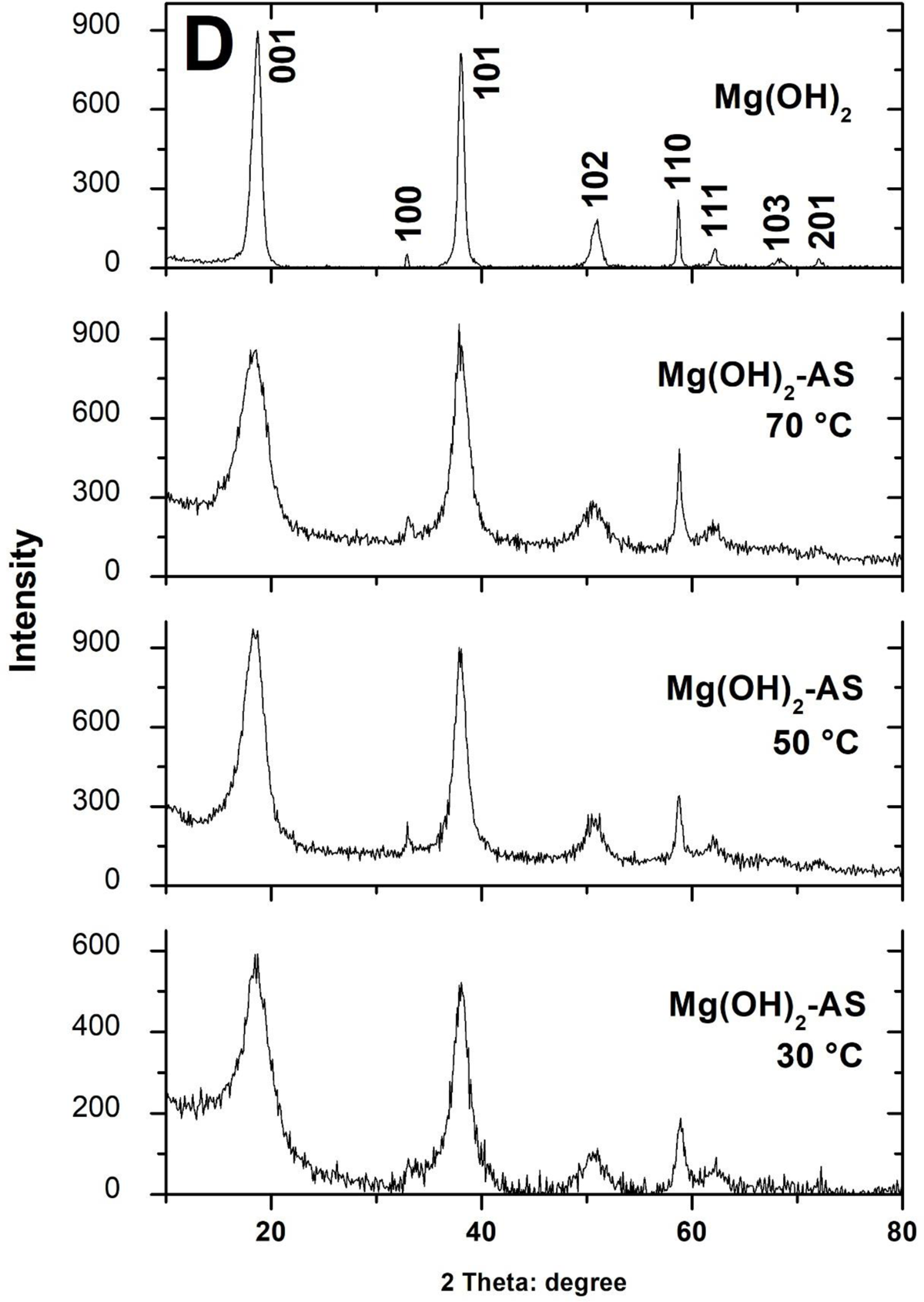
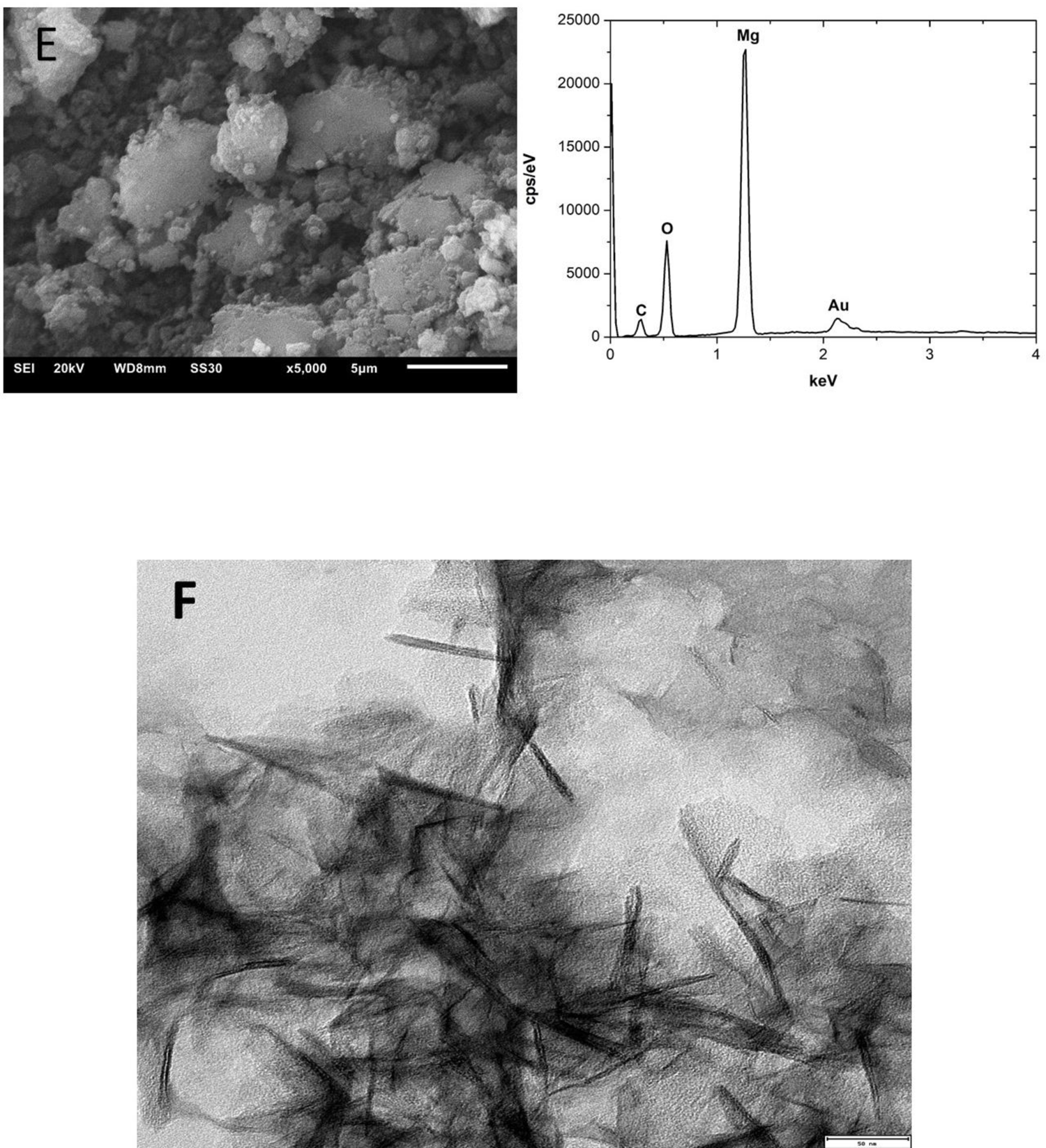
Physicochemical characterization of Mg(OH)_2_ nanoparticles prepared using AS extract. **A)** Visual observation of the colorations during the reaction of Mg^2+^ and AS extract at 30, 50 and 70 °C. **B)** Ultraviolet-visible spectra associated with the synthesis of Mg(OH)_2_-AS. Wavelengths of plasmon resonance peaks are labeled. **C)** Fourier transform attenuated total reflection infrared spectra of Mg(OH)_2_, AS and Mg(OH)_2_-AS (70°C). **D)** Powder X-ray diffractograms of Mg(OH)_2_ and Mg(OH)_2_-AS powders at 30, 50 and 70°C. **E)** Image of scanning electron microscopy and X-ray energy dispersive spectroscopy. **F)** image from transmission electron microscopy representing needles of 33±9 nm long and 4±2 nm wide considering 25 isolated needles, with scale bar depicting 50 nm.

### Cell morphology analysis

After 3 days of exposure with Mg(OH) _2_-AS or controls (basal media, Mg(OH)_2_ or AS, respectively), cells were washed with 500 μL PBS and permeabilized with 0.1 % (v/v) Triton-X 100 in PBS for 15 minutes at room temperature. After aspirating the solution, cells were incubated with 300 μL of a PBST blocking solution (0.1 % (v/v) Tween 20, 1% BSA in PBS) for 30 min at 4°C. Subsequently, cells were incubated with 100 μL of a Phalloidin-iFluor 488 working solution (1:500 dilution in PBST) over night at 4°C, protected from light. The next day, DAPI was added directly to the well to a final concentration of 5 μL/mL, and incubated for 10 minutes at 4°C. Afterwards, the staining solution was discarded, and each well was washed three times with 500 μL of PBS. Finally, samples were slide-mounted for microscopy (H-1000, Vectashield, Germany) and representative pictures were acquired using a fluorescence microscope (DMi8, Leica).

Automated quantification of cellular shape features was performed using the CellProfiler 4.2.5 pipeline, as previously described [34]. Only full segmented cells, i.e., single cells with non-overlapping borders and/or complete area of cell present within the borders of the image, were used for further analysis. The statistical significance of each morphological parameter (area, compactness, aspect ratio and solidity) was evaluated with the Kruskal-Wallis test against the control group in basal medium (p ≤ 0.05, GraphPad Prism 10.0.2). The parameters relevant for this analysis were: the area (counted pixels in the object defined by the segmentation, relatively scaled to µm^2^), the compactness (calculated as Perimeter2/4*π*Area, where a filled circle will have a compactness of 1, with irregular objects or objects with holes having a value greater than 1), the aspect ratio (the ratio of the minimum to the maximum feret diameter, giving an indication for the elongation of the object), and the solidity (as the proportion of the pixels in the convex hull that are also in the object).

### Osteogenic differentiation

To obtain preliminary data on the impact of Mg(OH)_2_-AS nanoparticles on BM-MSC osteogenic differentiation capacity, BM-MSCs of one donor were cultured for 3 weeks in osteogenic media in the presence of nanoparticles and respective controls, and the matrix-mineral deposition was assessed by Alizarin-red staining. BM-MSCs were seeded at 2 × 10^4^ cells per cm^2^ in 24-well plates in basal culture media. After confluence was reached, media was changed to i) complete osteogenic medium (DMEM low glucose medium supplemented with 10% FCS, 1% P/S, 1% HEPES, 50 µg/mL ascorbic acid-2-phosphate, 5 mM β-glycerophosphate and 10 nM dexamethasone (all Sigma-Aldrich)), ii) osteogenic medium without dexamethasone or iii) control medium without osteogenic factors ascorbic acid-2-phosphate, β-glycerophosphate and dexamethasone. The media was changed three times per week. Nanoparticles and control substances were subsequently added to the wells at a final concentration of 150 μg/mL and freshly supplemented upon each medium change. After 21 days of culture, wells were washed twice with cold PBS and fixed in 70% ethanol (Carl Roth) for 1 h at −20°C. After drying, deposited mineral was stained with a 2% Alizarin red solution (ScienCell, USA) for 15 min at room temperature under gentle agitation. After several washing steps, representative brightfield micrographs (DMi8, Leica) were taken. Afterwards, incorporated stain was quantified. For this, in 10% cetylpyridinium chloride solution (Sigma-Aldrich) was added to the wells for elution for 20 min under gentle agitation. Samples and a standard curve of diluted staining solution were measured at a microplate reader (infinite M200, Tecan, Switzerland) in a 96-well plate at 570 nm in duplicates. Results were calculated as µg/mL of Alizarin-red. Triplicates prepared from cells of one donor were analyzed for Mg(OH)_2_-AS experiments and presented as the mean ± standard error of the mean. Controls were made in duplicates. Data were compared by two-way ANOVA.

### Statistical analysis

Quantitative data was analyzed using Graphpad Prism for Windows software (version 9.1) and presented as the mean ± standard error of the mean. The results were compared by two-way ANOVA followed by Tukey’s multiple comparison at the 95% threshold.

## Results

### Plant extract studies

In order to extract the plant metabolites, 100 g of fresh *Anthocleista schweinfurthii* (AS) leaves were infused in 500 mL of distilled water, then dried and filtered. A brown powder with a mass of 1.6 g was obtained. Drying 100 g of fresh leaves in the open air and away from the sun produced a mass of 23.7 g of dry leaves, giving an extraction rate of 6.8%. The data are summarized in Table 1 below. Qualitative analysis of the aqueous extract of *Anthocleista schweinfurthii* revealed secondary metabolites such as alkaloids, flavonoids, coumarins, tannins, saponins and phenols (see Table 2).

**Table 1.** Extracted content and physicochemical characteristics of AS extract.

| Fresh leaves (g) | Dry leaves (g) | Dry extract (g) | Content (%) | Physicochemical characteristics |  |  |  |
| --- | --- | --- | --- | --- | --- | --- | --- |
| 100 | 23.7 | 1.6 | 6.8 | Color |  | Form |  |
|  |  |  |  | AE<br>Pale yellow<br>pH<br>5.3 | DE<br>Brown | AE<br>Liquid | DE<br>Powder |

AE: aqueous extract; DE: dry extract

**Table 2.**
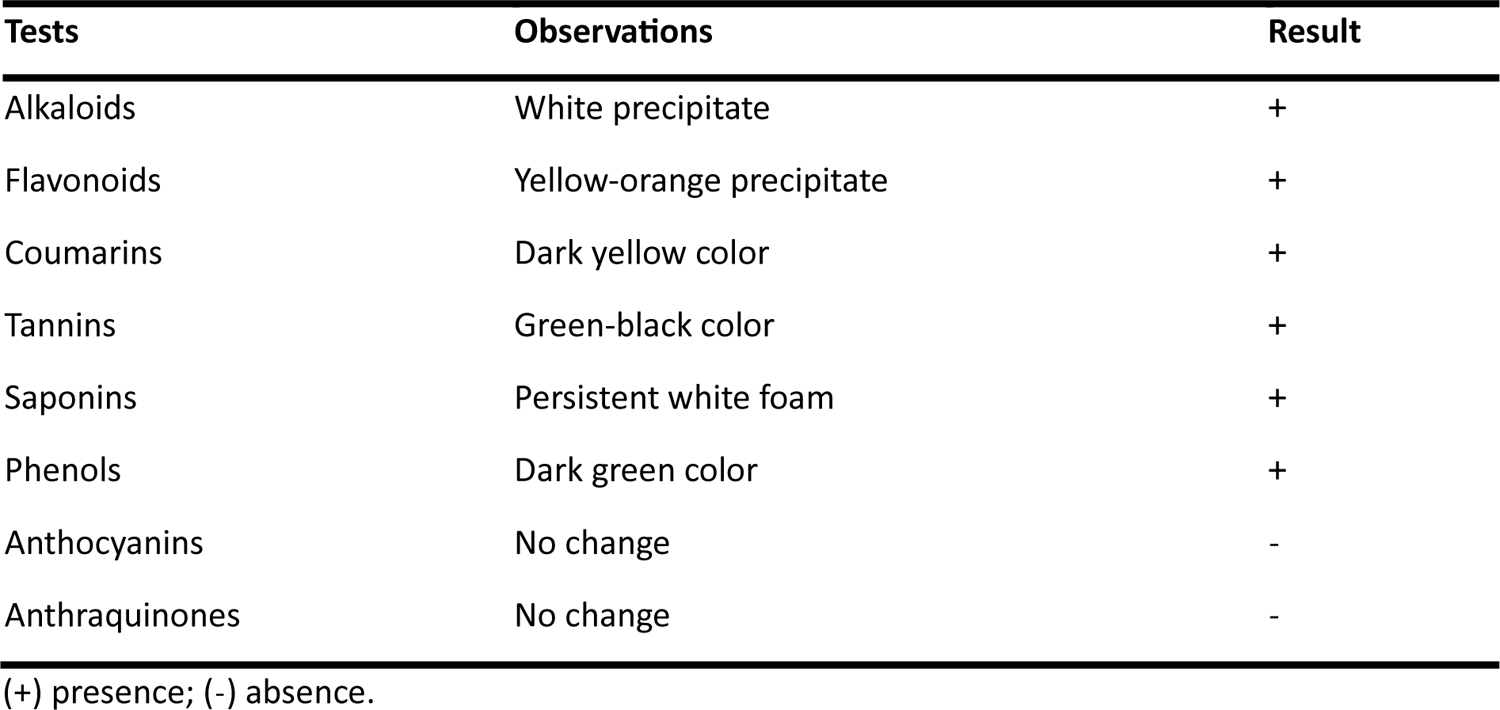
Phytochemical screening of AS extract.

### Nanoparticle analyses

Figure 2 shows the solution of magnesium nitrate salt, the *Anthocleista schweinfurthii* extract and of the nanoparticles obtained at 30, 50 and 70 °C. Visual observation of the hot reaction showed a yellowish coloration that becomes more consistent with increasing temperature. Experimental evidence for magnesium oxide nanoparticle formation was obtained by UV-visible spectroscopy (see Figure 2B). The curves after 2.5 hours of heating to 30, 50 and 70 °C showed an increase in absorbance intensity indicating the increase of nanoparticles density in the medium, with the maximum of the broad absorption moving to longer wavelengths with higher temperature due to increasing nanoparticles size. To determine the hydrodynamic radius of Mg(OH)_2_-AS at different temperatures, Dynamic Light Scattering (DLS) was used (Figure S1, SM). The particles were measured three times at each temperature and all particles show different size. The particles showed a distribution with polydispersing tendency at all temperatures and the hydrodynamic radius was between 271-3784 nm. The size of the particles increased at higher temperatures, likely because the stability of the particles decreases at higher temperatures, causing them to agglomerate more. We also applied Fourier transform infrared spectroscopy (Figure 2C) to study the metal interface composition. To achieve this aim Mg(OH)_2_, AS and Mg(OH)_2_-AS had to be measured separately. The superposition of the vibrations of AS, Mg(OH)_2_ and Mg(OH)_2_-AS show that the IR signature of Mg(OH)_2_ dominates in Mg(OH)_2_-AS, indicating that Mg(OH)_2_ is the major component of the composite. The C-H bands of AS below 3000 cm^-1^ were essentially no longer visible in the composite. The vibration at 3698 cm^-1^ is assigned to Mg-O [35,36]. The phase study by powder diffractometry obtained from grains at 30, 50, and 70 °C (Figure 2D) and the diffractogram of Mg(OH)_2_ nanoparticles indicated crystalline planes at positions (001), (100), (101), (102), (110), (111), (103) and (201) corresponding to Mg(OH)_2_ by comparison with The International Centre for Diffraction Data (ICDD) 44-1482 card. The nanoparticle sizes obtained by the Scherrer equation considering the highest intensity signal (101) are 53 nm at 30 °C, and 50 °C, 46 nm at 70 °C, and 8 nm for Mg(OH)_2_. SEM to study the morphology, shows spherical agglomerated particles (Figure 2E). Elemental characterization by EDX shows a composition of O, C and Mg, with little carbon content. Needles are obtained by transmission electron microscopy with mean sizes 33±9 nm long and 4±2 nm wide (Figure 2F).

### Acute cutaneous toxicity, wound healing and protein degradation

Acute toxicity experiment was conducted to study the animal growth during treatment by topical application and clinical parameters where recorded. During the acute toxicity assay, the animal growth and clinical parameters are not impaired by Mg(OH)_2_-whole AS extract nanoparticles compared to control rats who received water (Figure 3A and Table 3). The growth rate increases significatively at day 10 and 12 for animal treated with the nanoparticles suggesting that they are not affecting growing. The rates are similar again at day 14 suggesting that animals feed normally during the experiment as a sign of good health. The rat wound healing experiment by daily application of the nanoparticles to wounds revealed that the wound closure rate was significantly higher for animals treated with Mg(OH)_2_-AS 100 mg/kg nanoparticles when compared to whole AS extract, positive (trolamine) or negative (water) control groups (Figure 3B, C). Inflammation is the first stage in the wound healing process. To test whether Mg(OH)_2_-AS nanoparticles can affect inflammation, we used an *in vitro* assay, mixing the nanoparticle extraction with BSA [32]. We found that Mg(OH)_2_-AS inhibits heat induced protein degradation in a concentration-dependant manner and significantly higher compared to the non-steroidal anti-inflammatory drug diclofenac at 300 and 500 μg/mL (Figure 3D).

**Figure 3.**
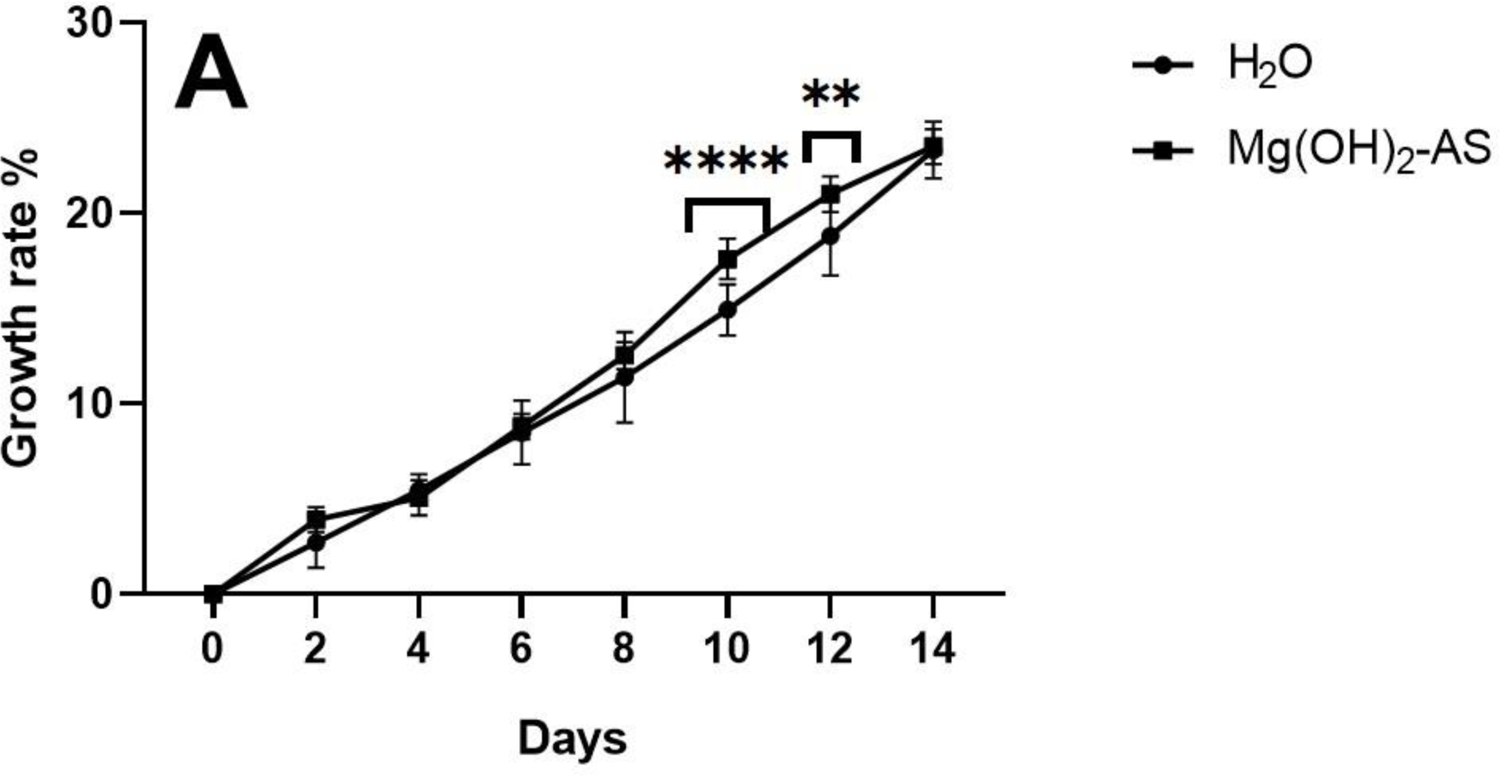
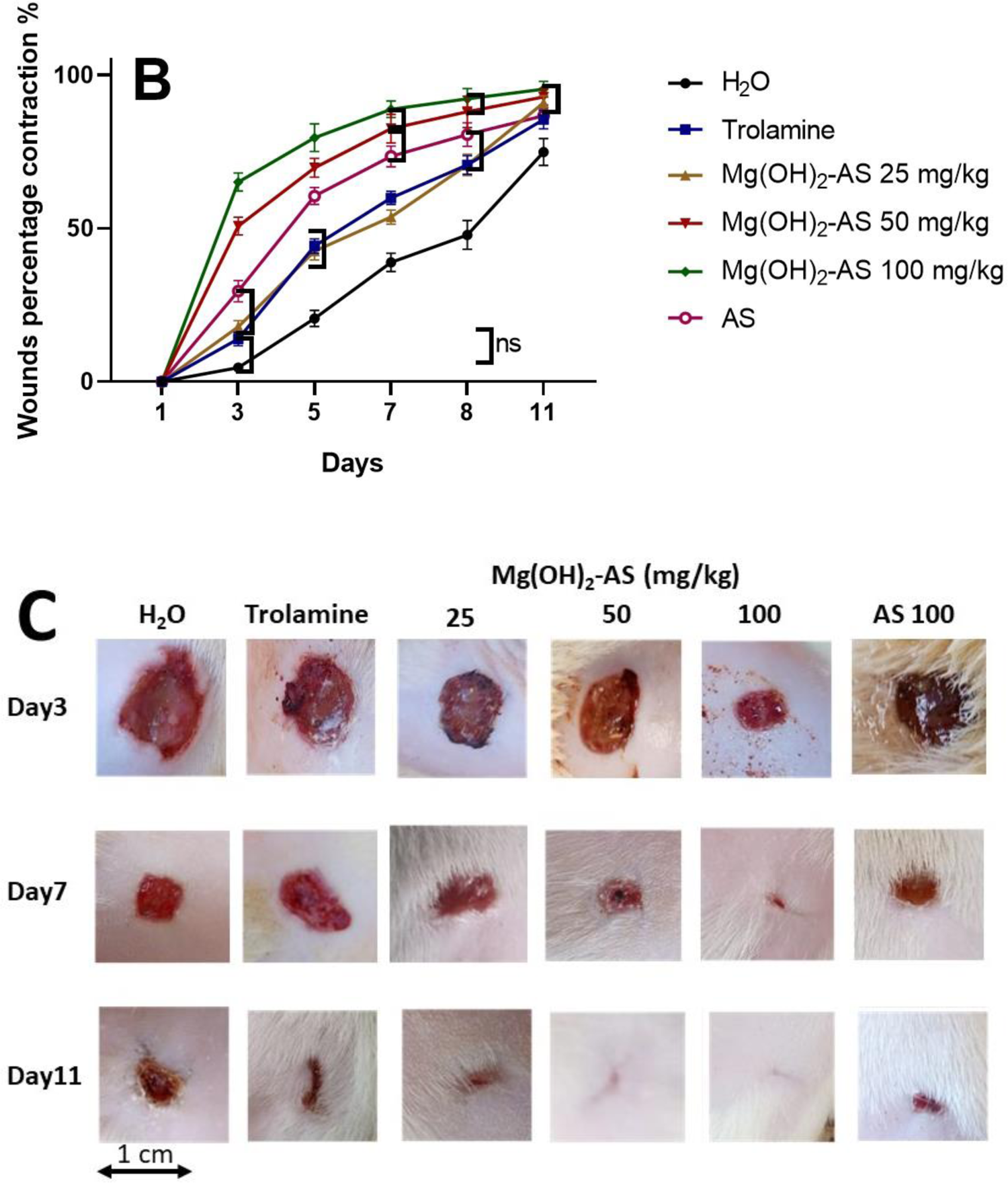
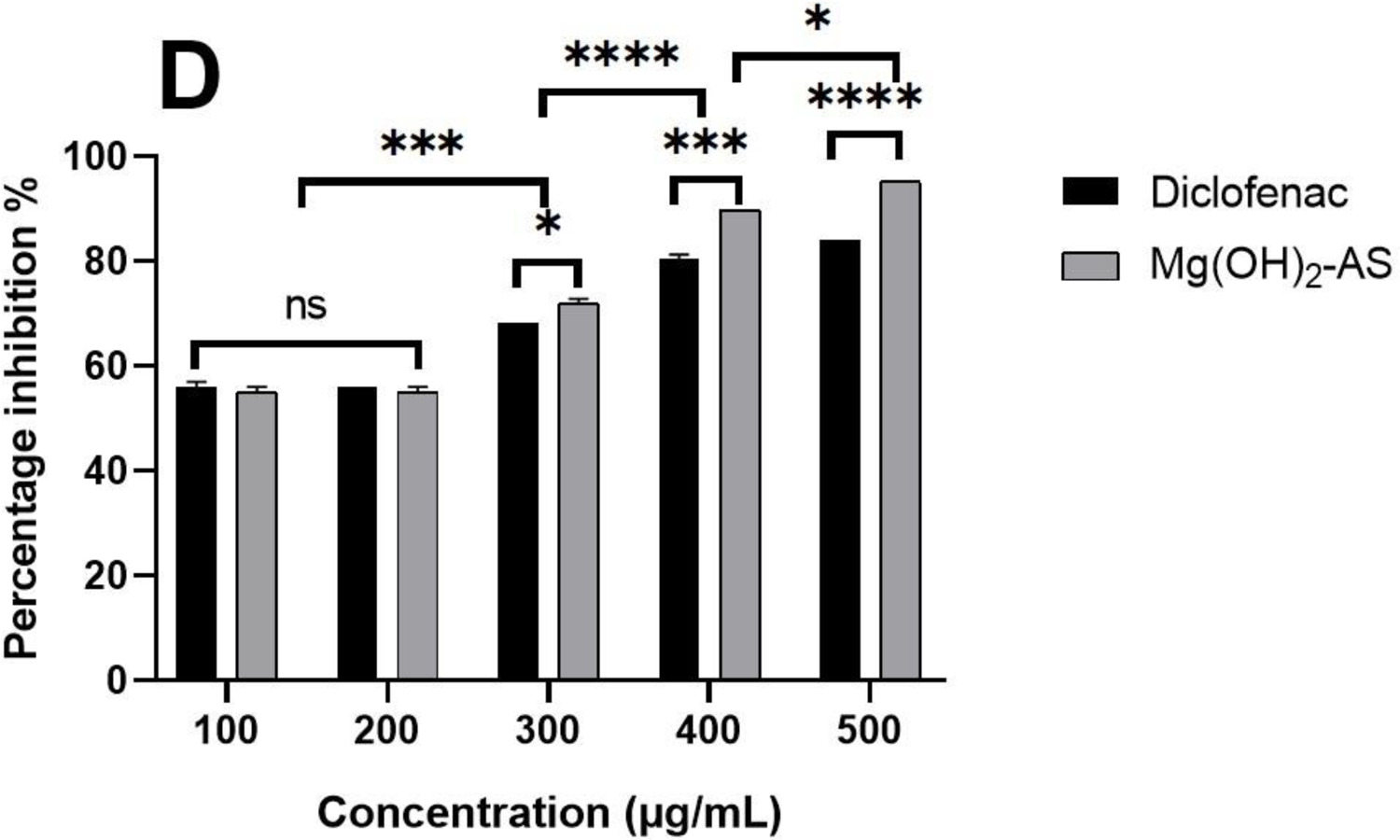
Wound healing and toxicity on Wistar rats. **A)** Growth rate curve (% of weight gain) of animals during the acute dermal toxicity assay. The mean of 3 measurement + standard error of the mean is shown. Two-way ANOVA was used for comparison. The weight evolution of rats subjected to the acute cutaneous toxicity test reveals a normal increase in rat weight, with significant difference between the control and test batches at day 9 and 11 (**** indicate p < 0.0001, *** p < 0.001, ** p < 0.01, * p < 0.05). **B)** Wound closure expressed as percentage contraction in animals at days 3, 5, 7, and 11 and **C)** representative pictures at day 3, 7 and 11, of wounds in animals treated with Mg(OH)_2_-AS (25, 50, 100 mg/kg), Trolamine, AS whole extract, or water. **D)** Inhibition of BSA degradation as an important anti-inflammatory cue by Mg(OH)_2_-AS nanoparticles and diclofenac. The mean of 3 measurement + standard error of the mean is shown. Two-way ANOVA was used for comparison.

**Table 3.**
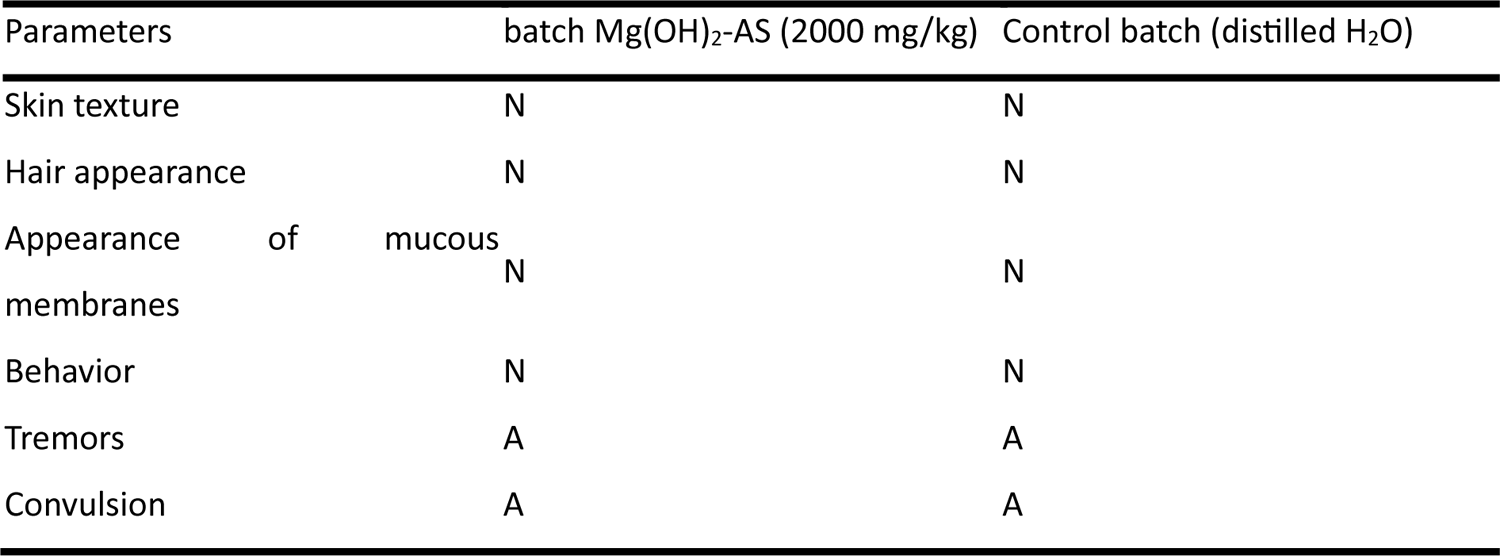
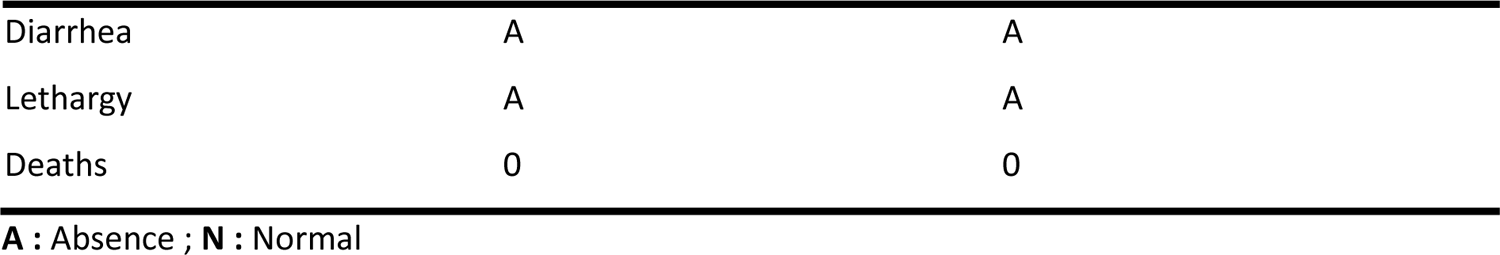
Monitoring of clinical parameters in rats during acute dermal toxicity test.

### Mesenchymal stromal cell metabolic activity, morphology and osteogenic differentiation

In order to evaluate the impact of nanoparticles on cell viability and proliferation, we investigated the behaviour of Mg(OH)_2_-AS nanoparticles on stromal cells as they are involved in tissue regeneration. bone marrow derived cells were selected to address particularly bone regeneration as model tissue for high regenerative capacities. Metabolic activity of bone marrow mesenchymal stromal cells (BM-MSCs) was monitored over 5 days. Whole AS extract and Mg(OH)_2_ treated cells showed a significantly reduced metabolic activity at all time points and both applied concentrations suggesting poor cell viability (Figure 4A). In contrast, Mg(OH)_2-_AS treated cells showed the trend of increased metabolic activity from day 1 to day 5, in particular at 300 μg/mL, suggesting that these nanoparticles might promote cell proliferation.

**Figure 4.**
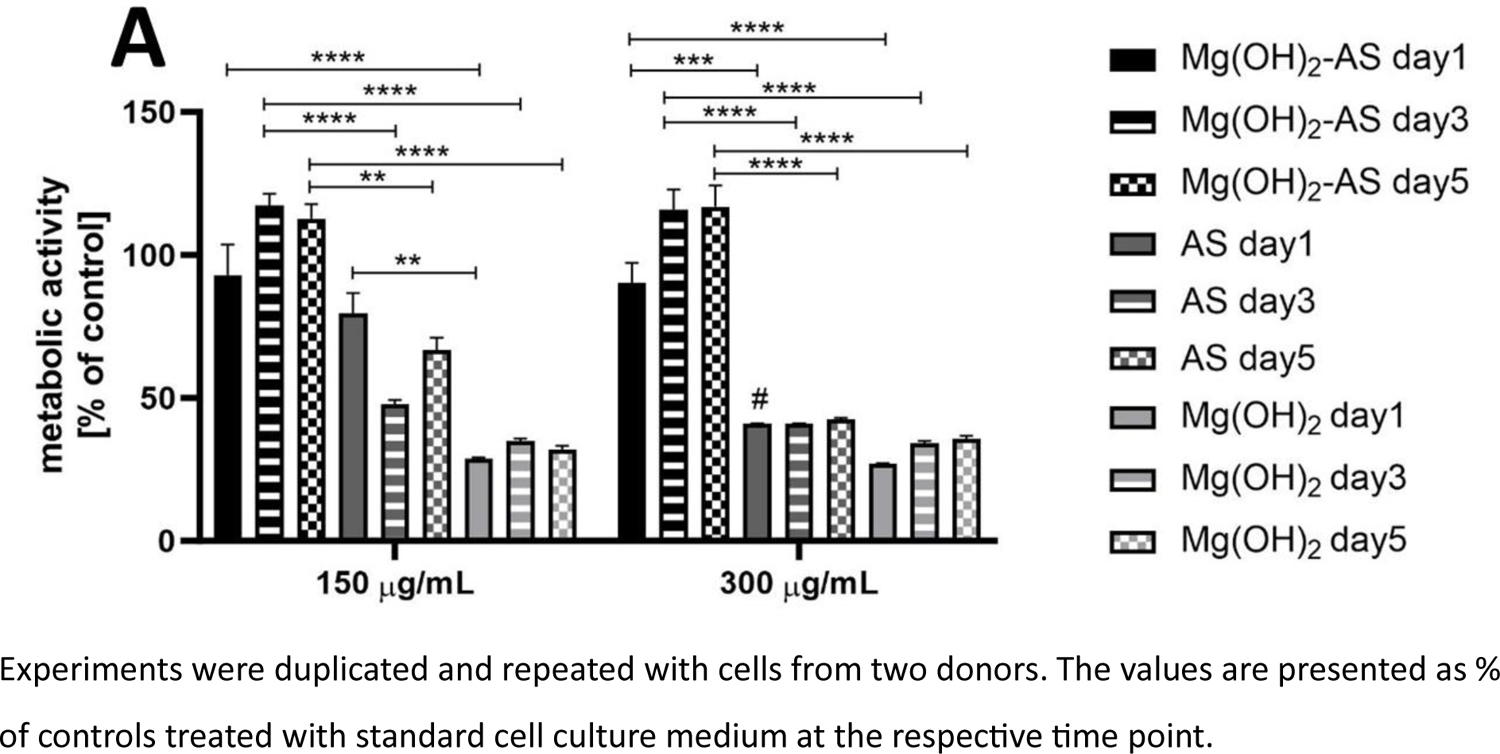
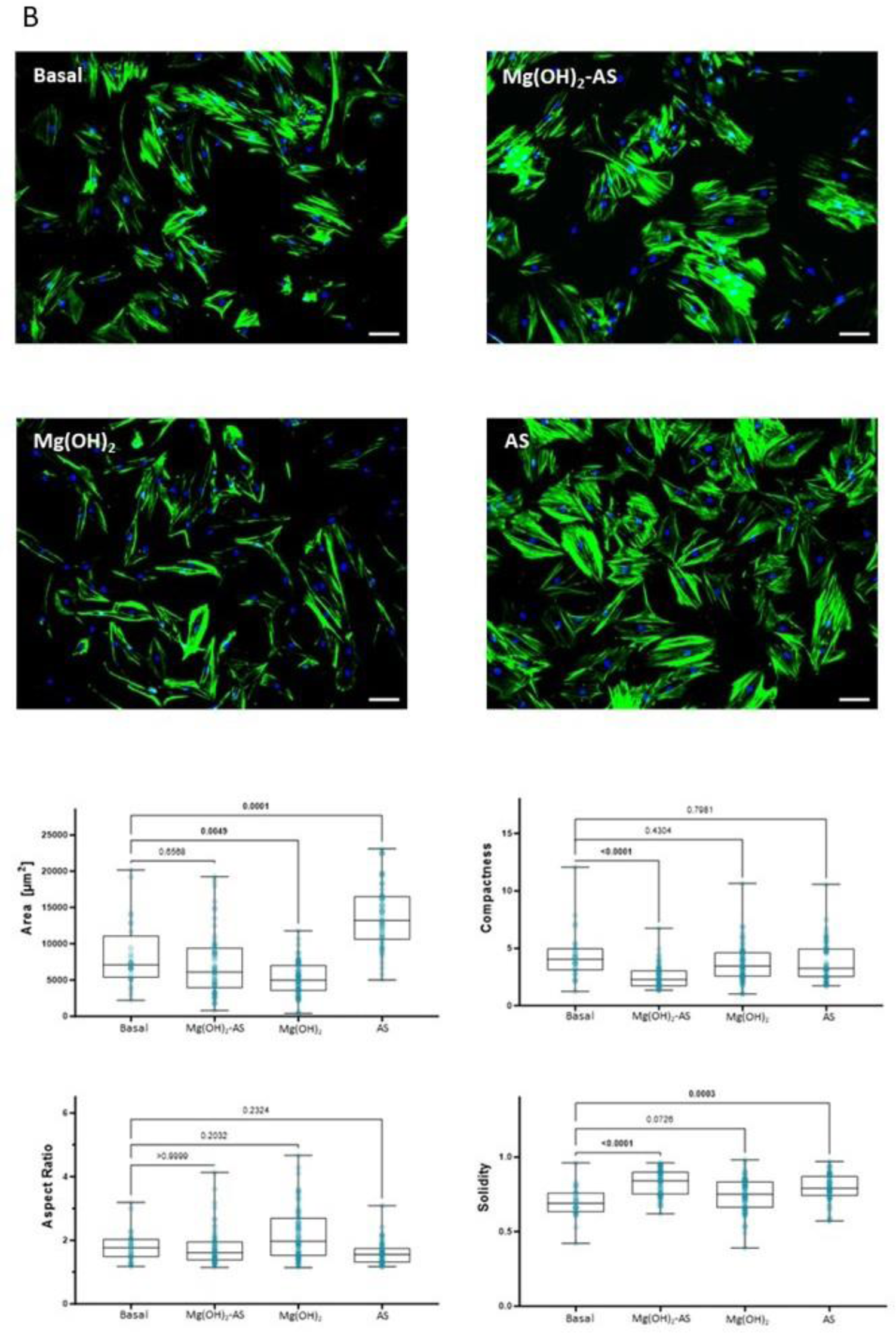
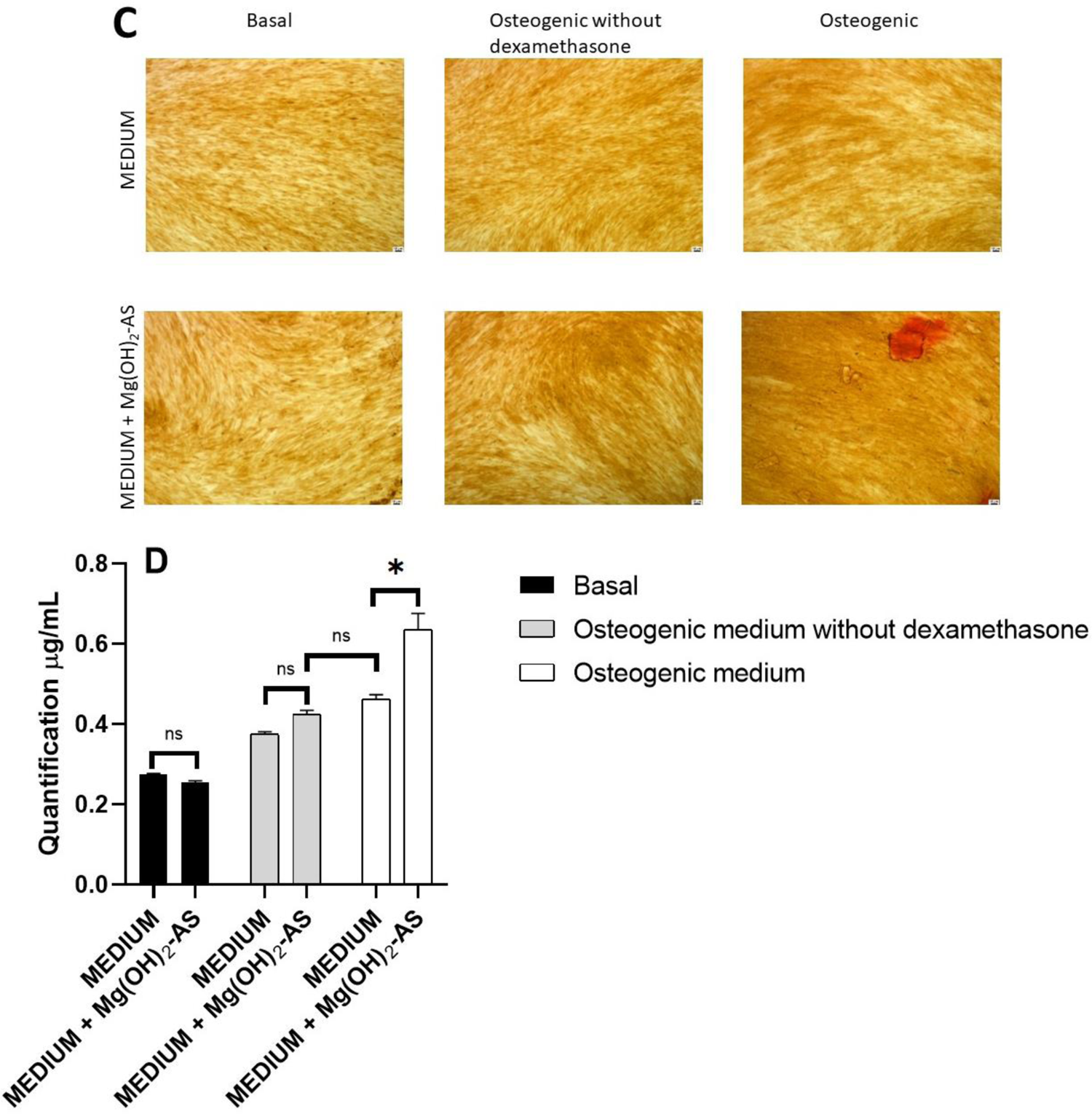
BM-MSCs metabolic activity, morphology and osteogenic differentiation **A)** Cell viability / proliferation (MTT assay) of MSCs cultured under basal conditions and exposed to Mg(OH)_2_-AS, AS and Mg(OH)_2_ nanoparticles at 150 μg/mL and 300 μg/mL, for 1, 3, and 5 days. n = 4 (duplicates were performed on cells from two donors) + standard error of the mean and two-way ANOVA is shown. **B)** Cell morphology analysis of MSCs cultured under basal conditions and treated with Mg(OH)_2_-AS nanoparticles, Mg(OH)_2_ nanoparticles and AS 150 μg/mL for 3 days. On top, representative composite images of cells stained with Phalloidin-iFluor 488 conjugate to stain actin filaments within the cytoplasm (green) and DAPI to stain the nucleus (blue), scale bar 100 µm. Bellow, analysis of single-cell morphological descriptors (area, compactness, aspect ratio and solidity) exported from CellProfiler pipeline (Basal: n=28, Mg(OH)_2_-AS: n=57, Mg(OH)_2_: n=69, AS: n=54 segmented cells). Quantitative differences presented as their normal distribution, with mean value and Min-to-Max bars, per culture condition with statistical significance set to p ≤ 0.05, in bold (Kruskal-Wallis, multiple comparison method). **C)** Representative images of Alizarin Red S staining at day 21 of basal, and osteogenic culture. Scale bar: 100 μm. **D)** Osteogenic potential assessed by quantification of incorporated Alizarin Red S dye at day 21. n = 3 (Triplicates of cells from one donor) + standard error of the mean and two-way ANOVA is shown.

Interestingly, when we performed single cell morphological analysis obtained from fluorescence microscopy, we observed adherent cells with typical fibroblastic morphology in both control (basal medium) and Mg(OH)_2_-AS treated cultures. However, cells cultured with Mg(OH)_2_-AS seem to be significantly more compact and with less irregular borders (i.e., higher solidity), demonstrating favorable conditions of the substrate for cell culture. On the other hand, treatment with Mg(OH)_2_ alone seems to be not really favorable, showing considerably smaller cells (illustrated by an untypical spreading) than in the basal condition, confirming the metabolic activity results. Finally AS-treated cells acquired a significant big size and round shape, typical of senescent cells and therefore deviating from the osteogenic differentiation lineage [25].

To obtain a first insight into the impact of Mg(OH)_2_-AS nanoparticles on osteogenic differentiation of BM-MSCs, we exposed BM-MSCs from one donor to either control or osteogenic medium for 21 days and assessed brown to red staining of deposited mineral in differentiated cells (Figure B). We quantified a significant increase in incorporated Alizarin Red stain in the Mg(OH)_2_-AS nanoparticle treated group, compared to standard osteogenic medium without supplementation, indicating more mineral deposition and thus a stronger differentiation (Figure C).

Experiments were duplicated and repeated with cells from two donors. The values are presented as % of controls treated with standard cell culture medium at the respective time point.

## Discussion

Magnesium hydroxide from plant extracts can be good candidates in injury recovering and in bone regenerative medicine by enhancing biocompatibility, cell proliferation and potentially osteogenic differentiation. The nanoparticle synthesis required an aqueous extract. Our measurement of 6.8% extractable content for *Anthocleista schweinfurthii* is comparable to that reported by Tchangou et al., 6.9%, on *Stryvhnos phaeotricha*, a plant of the same family [37]. Phytochemical screening of AS revealed secondary metabolites such as alkaloids, flavonoids, coumarins, tannins, saponins and phenols, and the absence of terpenes, anthocyanins and anthraquinones. It should be noted that previous phytochemical screening studies of a methanolic extract of the leaves of this plant revealed the presence of polyphenols, alkaloids, terpenes and steroids (DRC), tannins and phenols (West and South West Cameroon). These results underline the diversity of chemical compounds present in AS leaves collected for this study in the littoral and could highlight the importance of variation in phytochemical profiles according to environmental and geographical factors, and the extraction methods used [38,39]. The synthesis of Mg(OH)_2_ nanoparticles in this work required heat to increase contact between Mg^2+^ ions and extract metabolites. In addition, the use of OH^-^ ions is known to have a positive effect on synthesis by reacting with H^+^ ions in carboxylic acids and phenols, making them more reactive. The shift of surface plasmon resonance observed by UV-Vis spectroscopy from left to right (pure Mg(OH)_2_ to plant mediated magnesium nanoparticles), can be explained by the increase in size after plant metabolites reduction. The increase in coloration from 30 to 70 °C goes hand in hand with the increase in absorbance, both of which are linked to the rise in nanoparticle solution density [40]. This observation is corroborated by DLS measurements of intensity, volume and number, which show a rapid evolution of dispersion during different scans. This rapid evolution may be related to the use of water, a solvent with a small partial charge that causes more attractive interactions than repulsion interactions, situation found in different types of dispersed nanomaterials, including nanosized plants [41]. The polydispersity of hydrodynamic diameters decreases with time, which can be related to the reduction of agglomeration. The diffractograms show a pattern with noise in the background corresponding to a decrease in crystallinity. Pinho et al (2021ab) have obtained a rose hip functionalized magnesium hydroxide consisting of amorphous nanoparticles. Crystallinity seems to be better when the precursor is a nitrate [35,36]. The sizes obtained by the Scherrer relation are below 60 nm for the three temperatures used 30, 50 and 70 °C. These size are larger to 8 nm obtained for pure Mg(OH)_2_ particles and supports the UV-Vis analysis. An agglomerated form is detected by SEM with elemental analysis showing the presence of C, O and Mg ions, proving both an interface bearing AS metabolites and the presence of Mg(OH)_2_ hydroxide. TEM shows that Mg(OH)_2_ nanoparticles present needle-like shapes, while other shapes such as sheets and squares or spheres have been obtained by synthesis using plants [35,36].

Importantly, no signs of toxicity were observed during the acute dermal safety test. The lethal dose 50 (LD_50_) of these nanoparticles is greater than 2000 mg/kg. Mg(OH)_2_-AS nanoparticles are therefore not classified in the OECD category of products at risk of acute cutaneous toxicity. In addition, Mg(OH)_2_-AS did not affect the weight gain of the animals. Body mass time points are significatively increasing, a situation observed with silver metallic nanoparticles driving metabolites and encapsulated by chitosan [37]. The development of novel, effective, low-toxicity methods is fundamental for their use as treatments.

In animal models with wounds induced, wound treated with Mg(OH)_2_-AS healed faster than those treated with the positive control. A similar progression was observed with biogenic silver nanoparticles [30]. Magnesium nanoparticles can promote wound healing by releasing magnesium ions that stimulate the proliferation of fibroblasts, a population of cells responsible for matrix synthesis and tissue repair [42]. Our results from the protein degradation assay also suggest that Mg(OH)_2_-AS could reduce inflammation, which would favour an environment conducive to healing by reducing oxidative stress and promoting tissue regeneration. Chronic inflammation is often seen in inflammatory arthritis, chronic osteomyelitis, non-union of fractures, and osteonecrosis [43]. Thus, an anti-inflammatory characteristic of nanoparticles is desirable for new therapeutic strategies targeting such diseases.

Here, we studied cellular behaviors upon treatment with 150 and 300 ug/mL, based on previous studies showing that 300-500 ug/mL of MgO nanoparticles kills 50% of BM-MSC cells and rosehip-functionalized MgO nanoparticles did not affect cell viability and proliferation in the concentration range of 1 to 100 ug/mL [35,36,44]. We showed that cells treated with Mg(OH)_2_-AS showed higher metabolic activity compared to cells treated with Mg(OH)_2_ or AS alone. The cells are able to grow at 300 μg/mL Mg(OH)_2_-AS, but cell death was observed with AS and Mg(OH)_2_. Viability using Mg(OH)_2_-AS was observed at 300 μg/mL in this research. Proliferation trended towards increasing with time for Mg(OH)_2_-AS, in contrast to whole AS extract and pure Mg(OH)_2_, which decreased cell density. This behaviour of BM-MSCs is in line to the results reported with MgO by Wetteland et al., 2016 who obtained smaller cells with signs of membrane damage [44].

Nanoparticle bioactivities are related to physical properties of the metal and their mode of action. Thus, nanoparticles toxicity could be due to direct contact with the oxide in the culture medium [44]. Our study study shows that synthesized Mg(OH)_2_ and AS have no beneficial activity on BM-MSCs in the MTT assay. Contact of BM-MSCs with metabolites on the surface of Mg(OH)_2_-AS seems to have a more beneficial effect than direct contact with Mg(OH)_2_. These observations are corroborated by the observation of healthy, abundant, confluent cells for Mg(OH)_2_-AS at 150 μg/mL comparable to the control by fluorescence microscopy. Based on the fluorescence images, the treatment condition with AS seems not to impact cell density; however the cells acquire a significant big size and round shape, which is typical for senescent cells [25]. On the other hand, the treatment with Mg(OH)_2_ seems to be unfavourable for the cells; they are considerably smaller than the basal condition and not very viable according to the MTT results. The combination with Mg(OH)_2_-AS seems to achieve a good balance, where the cell size is not significantly different from the basal, however, cells seem to be significantly more compact and with less irregular borders, demonstrating favourable conditions of the particles for cell culture. At the studied timepoint, the aspect ratio, which is known to be an early predictor of osteogenic commitment [45], does not seem to be changed significantly in any condition. One limitation of this analysis is that is the quantifications are based on one representative image for each condition. However, the observed differences are in line with the data from MTT analysis. The preliminary osteogenic differentiation experiment shows abundant brown staining of BM-MSCs treated with Mg(OH)_2_-AS nanoparticles in dexamethasone-supplemented osteogenic medium. Based on the quantification of the incorporated dye, the latter is significant compared to the basal media indicating a stonger differentiation.

This positive activity may be due to the release of Mg^2+^ ions from Mg(OH)_2_-AS known to enhance osteogenic activity of BM-MSCs [46]. Spherical magnesium hydroxide mediated rose hip and Mg(OH)_2_ with osteoblastic properties have been shown to be active on the MG-63 cell line [35]. Pinho et al. postulated a beneficial impact of polyphenols in rosehip extract polyphenols. The expression of early osteogenic differentiation markers such as the transcription factor Runx2, type I collagen and alkaline phosphatase, as well as later differentiation markers such as Osterix (SP7) and osteopontin (SPP1), was higher in cultures exposed to synthesized Mg(OH)_2_ rose hip derived nanoparticles [35]. The Mg(OH)_2_-AS nanoparticles structure with an interface covered with plant secondary metabolites prevents direct contact with cells, resulting in low toxicity. An improved biocompatibility is generally achieved by interface modification using coating agents [47]. Alternatively, or in addition, there may be synergy with magnesium oxide acting as a transporter of metabolites to MSCs. Other nanoparticles, such as hydroxyapatite nanoparticles, silica nanoparticles and calcium carbonate nanoparticles, have been shown to have a positive role in promoting osteogenesis in MSCs [7]. This experiment suggest a positive impact on BM-MSC osteogenic differentiation. As study limitations, we must note that for experiments with primary human cells the number of repetitions with different donors was limited and thus variability of results in different donors could not be assessed at this point but will be the subject of future experiments.

## Conclusion

This work aimed to develop green Mg(OH)_2_ nanoparticles as materials for the management of injuries with bone fractures. Mg(OH)_2_-AS nanoparticles were obtained by treatment of magnesium nitrate with an aqueous extract of *Anthocleista schweinfurthii* in a basic medium and at temperatures up to 70 °C. These nanoparticles rapidly polydisperse in aqueous media and appear as aggregate and needles by electron microscopy. No acute dermal toxicity of Mg(OH)_2_-AS nanoparticles was found. In animal models, we observed high wound closure and high potential to prevent protein degradation as an anti-inflammatory cue. We also found that treatment with Mg(OH)_2_-AS is beneficial for the proliferation of mesenchymal stromal cells and seems to enhance osteogenic differentiation in media supplemented with ascorbic acid, β-glycerophosphate and dexamethasone. This study motivates further research into the potential use of Mg(OH)_2_-AS in the treatment of musculoskeletal disorders that have potential clinical applications for the repair of damaged or diseased tissue.

## Ethics Statement

The animals were examined and adapted to the new environmental conditions for a week before the formal experiment. All experimental procedures were in strict compliance with the approved protocol by the Institutional Ethic Committee For Human Research of the University of Douala (Protocol approval number 3552CEI-UDo/08/2023/T). Human bone marrow material was obtained as surgical waste material after approval of the local ethical board (University Hospital Wuerzburg, approval number 187/18).

## Acknowledgements

FEM thank the DAAD for a generous Visiting Professor Fellowship (grant no. 57588364). Support of the Imaging Core Facility, Biocenter, University of Würzburg, Germany for the JEOL JEM-2100 Transmission Electron Microscope funded by the Deutsche Forschungsgemeinschaft (DFG, German Research Foundation) - 218894163 is acknowledged. FEM received a Justice, Equity, Diversity and Inclusion Award from the Life Science Editors Foundation for editing support.

## Data availability

The authors declare that the data supporting the findings of this study including raw data files are available from the corresponding author upon reasonable request.

