## Supplemental 1 for "Magnesium hydroxide nanoneedles derived from *Anthocleista schweinfurthii* Gilg (Loganiaceae) support mesenchymal stromal cell proliferation and wound healing"

Supplement 1 : Dynamic Light Scattering

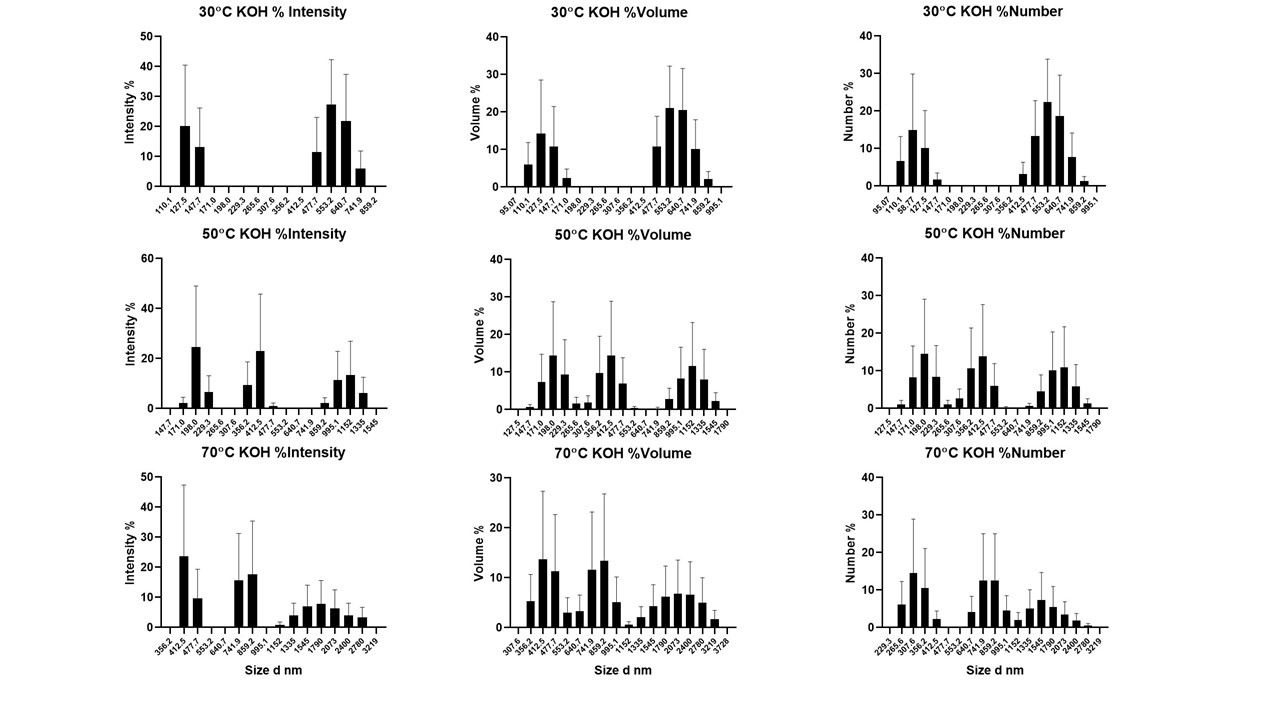

Table 1 : Assay

| Mg(OH)_2_-AS | Z-Ave | PdI |
| --- | --- | --- |
| T °C/Run | r.nm |  |
| 30/1 | 926 | 1 |
| 30/2 | 1471 | 0,976 |
| 30/3 | 1804 | 0,147 |
| 50/1 | 825,5 | 1 |
| 50/2 | 1240 | 0,972 |
| 50/3 | 1703 | 0,469 |
| 70/1 | 1678 | 0,72 |
| 70/2 | 2260 | 0,531 |
| 70/3 | 2435 | 0,259 |

Table 2 : Summary

|  | Temperature | Size (nm) | PdI |
| --- | --- | --- | --- |
| Mean | 20 | 1594 | 0,675 |
| Std Dev | 0.0 | 546,7 | 0,337 |
| RSD% | 0 | 34.3 | 49.9 |
| Minimum | 20 | 825.5 | 0.147 |
| Maximum | 20 | 2435 | 1.000 |

RSD Relative Standard Deviation
